# Dysregulation of NRSF/REST via EHMT1 is associated with psychiatric disorders

**DOI:** 10.1101/2021.04.26.441439

**Authors:** Mouhamed Alsaqati, Brittany A Davis, Jamie Wood, Megan Jones, Lora Jones, Aishah Westwood, Olena Petter, Anthony R Isles, David Linden, Marianne Van den Bree, Michael Owen, Jeremy Hall, Adrian J Harwood

## Abstract

Genetic evidence indicates disrupted epigenetic regulation as a major risk factor for psychiatric disorders, but the molecular mechanisms that drive this association are undetermined. EHMT1 is an epigenetic repressor that is causal for Kleefstra Syndrome (KS), a neurodevelopmental disorder (NDD) leading to ID, and is associated with schizophrenia. Here, we show that reduced EHMT1 activity decreases NRSF/REST protein leading to abnormal neuronal gene expression and progression of neurodevelopment in human iPSC. We further show that EHMT1 regulates NRSF/REST indirectly via repression of miRNA leading to aberrant neuronal gene regulation and neurodevelopment timing. Expression of a NRSF/REST mRNA that lacks the miRNA-binding sites restores neuronal gene regulation to EHMT1 deficient cells. Importantly, the EHMT1-regulated miRNA gene set with elevated expression is enriched for NRSF/REST regulators with an association for ID and schizophrenia. This reveals a molecular interaction between H3K9 dimethylation and NSRF/REST contributing to the aetiology of psychiatric disorders.

## Introduction

Genetic evidence points to altered epigenetic regulation as a substantial risk factor for many common psychiatric disorders (Lasalle, 2013; Kuehner et al., 2019). Genome-wide association studies (GWAS) have revealed multiple risk loci for neurodevelopmental disorders (NDD) that are associated with genes encoding epigenetic regulators. These often encompass more than one condition, including Intellectual Disability (ID), autism spectrum disorders (ASD) and schizophrenia. Accordingly, alleles affecting epigenetic regulatory mechanisms are associated with a range of psychiatric symptoms, including cognitive deficits, autistic traits and psychosis. In particular, epigenetic-related risk alleles are linked with biological pathways that converge on chromatin regulation via control of nucleosome positioning and histone methylation leading to altered gene transcription (De Rubeis et al., 2014; Gusev et al., 2018).

Studies on rare, often more penetrant, loss of function (LOF) gene variants associated with NDD reinforce this view. Disruption of the SNF-2 family chromatin re-modellers CHD7 and CHD8 are strongly associated with ID and along with CHD2 confer risk for ASD (Moccia and Martin, 2018). Histone lysine methyltransferases (KMT) are key epigenetic regulators and many are associated with NDD and psychiatric disorders. Loss of function (LOF) mutations of MLL3 (KMT2C), MLL5 (KMT2E), ASH1L (KMT2H), SUV420H1 (KMT5B) and histone lysine demethylases (KDM), KDM5B and KDM6B are all associated with ASD (Faundes et al., 2018; Lebrun et al., 2018; Stolerman et al., 2019). LOF variants of the H3K4 methyltransferase SETD1A are associated with schizophrenia (Singh et al., 2016). Severe neurodevelopmental disruptions and ID associated with mutation of a KMT underlie the genetic syndromes of Wiedemann-Steiner Syndrome (KMT2A), Kabuki Syndrome (KMT2D) and Kleefstra syndrome (EHMT1 or KMT1D). The latter is also associated with autistic features, psychosis and schizophrenia (Kirov et al., 2012; Adam and Isles, 2017). Although the genetic case for epigenetic regulation is thus well established, there is little knowledge of the downstream molecular mechanisms that link their actions to the underlying pathophysiology of the NDD. To address this question, we have investigated the mechanism by which reduced EHMT1 activity leads to an altered neurodevelopmental programme in both isogenic cell models of Kleefstra syndrome and patient-derived iPSC.

Kleefstra syndrome (KS) arises from a sub-telomeric microdeletion at 9q34, resulting in a heterozygous deletion of approximately ~700-kb (Kleefstra et al., 2006). This region contains at least five genes, including *ZMYND19, ARRDC1, C9ORF37, EHMT1*, and *CACNA1B* (Willemsen et al., 2012)’ (Kleefstra et al., 2006), however, the core clinical phenotypes are driven by haploinsufficiency of *EHMT1* (Kleefstra et al., 2006). EHMT1 is the primary enzyme for dimethylation of histone H3 at Lys9 residues (H3K9me2) (Shinkai and Tachibana, 2011) and is generally associated with transcriptional gene silencing (Tachibana et al., 2005). Genetic manipulation of EHMT1 has been studied in *Drosophila* and in rodent models at molecular, cellular and behavioural levels. *Drosophila ehmt1* mutants exhibit decreased dendrite branching of sensory neurons and impaired short and long-term memory that is reversed by restoring EHMT expression (Kramer et al., 2011). *EHMT1*^-/+^ mice demonstrate cranial abnormalities, hypotonia and delayed postnatal growth (Balemans et al., 2014). Functionally they show deficits in fear extinction and novel and spatial object recognition (Balemans et al., 2014; Sharma et al., 2017; Benevento et al., 2016; Davis et al., 2020). In rodent primary neuron cultures EHMT1 regulates the dynamics of multiple neural processes, including synaptic scaling and responses to addiction and stress (Benevento et al., 2016; Covington et al., 2011). To date, studies on KS patient-derived iPSC have focussed on differentiated, mature neurons, and show altered neuronal activity, synaptic signalling and network properties (Nagy et al., 2017). This demonstrates an effect of the 9q34.3 deletion on neuronal function, here we identify a molecular biological pathway that is disrupted in cellular models of KS.

We identify a EHMT1-mediated regulation of the transcriptional repressor NRSF/REST, a neuron-specific genes regulator (Hwang et al., 2017; Qureshi and Mehler, 2009; Schoenherr and Anderson, 1995). When bound to DNA, NRSF/REST acts as an epigenetic regulator by recruitment of a range of histone modifiers, including histone deacetylases (HDACs), SNF2-family chromatin remodellers and the EHMT1 paralogue, EHMT2 (KMT1C) (Ballas et al., 2005; Roopra et al., 2004). NRSF/REST in turn is regulated via microRNA (miRNA) that target the 3’UTR of its mRNA (Wu and Xie, 2006), and we show that EHMT1 suppresses expression of brain-related miRNAs including miR-153, miR-26a and miR-142. Examination of the altered miRNA expression profile in relationship to existing GWAS studies indicates a broader association between EHMT1 regulated miRNA, NRSF/REST and both ID and schizophrenia. Loss of *EHMT1* gene activity leads to elevated neuronal gene expression and can be reversed by re-expression of a *NRSF/REST* gene that lack the miRNA-regulatory sites. We further show that these gene regulatory abnormalities have substantial effects on *in vitro* neurodevelopment, causing premature neurodifferentiation and neuronal dysfunction.

## Results

### Loss of Ehmt1 reduces expression of NRSF/REST and increases expression of REST-target genes

We examined the effect of *Ehmt1* hemizygosity on the expression of neurodevelopment-specific genes in a mouse embryonic stem cell (mESC) model. mESC *Ehmt1*^+/-^ and *Ehmt*^flp^ control cell lines were cultured to neural progenitor cell stage (NPC) and gene expression examined using qRT-PCR (Figure 1A), (Table S1). Expression of *Rest* mRNA was decreased more than 6-fold and 10 out of 13 Nrsf/Rest-regulated genes in the study showed a significant increase in gene expression at the NPC stage. Based on these data we proposed that Ehmt1 may regulate neuronal gene transcription via control of Nrsf/Rest. To pursue this hypothesis, we treated wild type mESC with UNC0638, a highly potent and selective inhibitor of EHMT histone methyltransferases (Vedadi et al., 2011). As NRSF/REST is expressed in pluripotent stem cells, where it is suppresses neuronal gene expression, we tested mESC in the pluripotent state by treatment for 72 hours with a range of UNC0638 dosages. Western blot analysis showed a dose-dependent decrease of H3K9me2, with a half-maximal change at 200 nM (Figure 1B). This was accompanied by a similar dose-dependent decrease in NRSF/REST protein in cells treated with UNC0638 relative to the control (Figure 1B).

**Figure 1:**
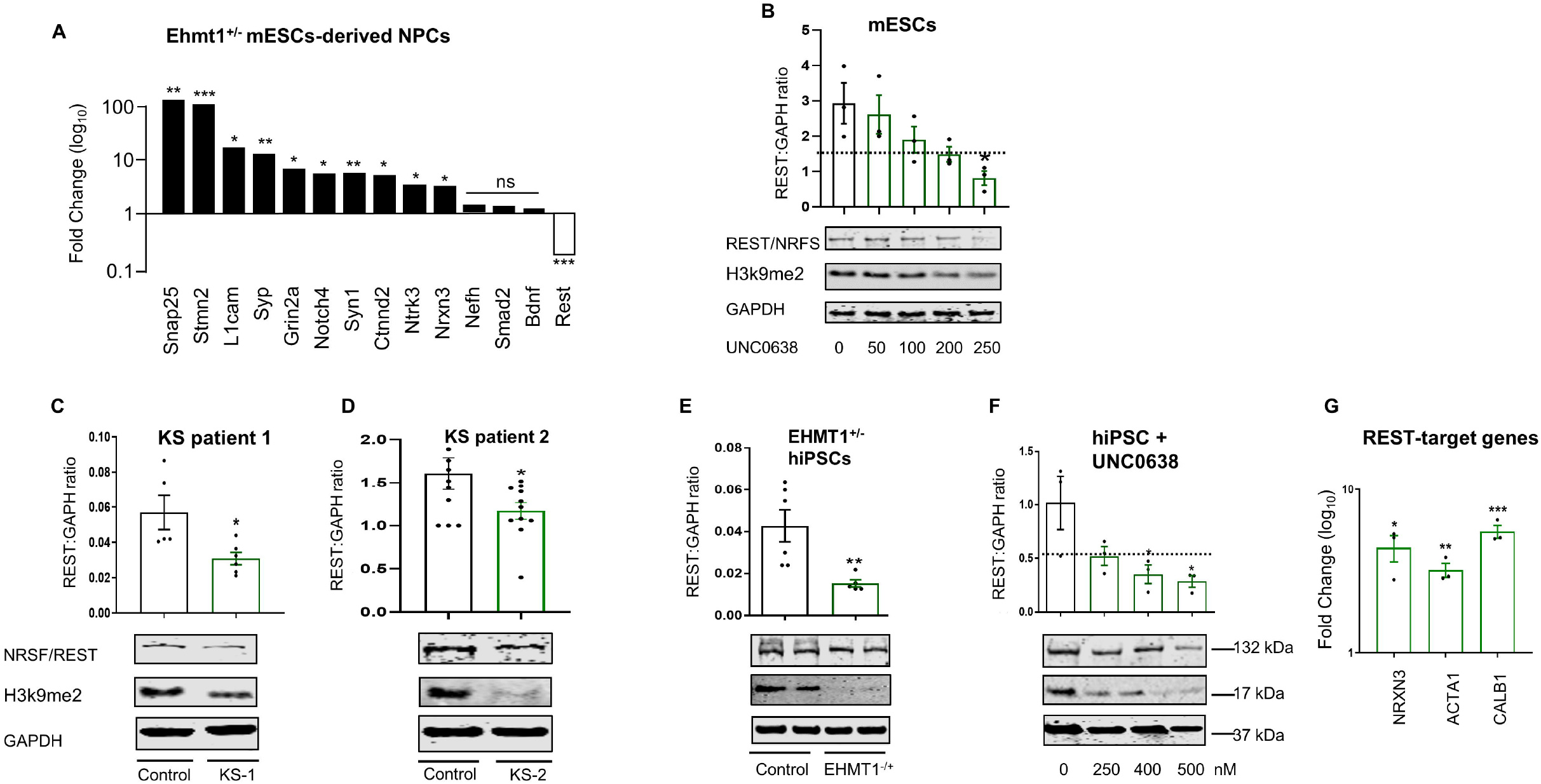
Expression of NRSF/REST and its target-genes in reduced EHMT1 activity and KS patient iPSC. (A) qRT-PCR analysis of neuron-specific genes in Ehmt1^-/+^ mESCs-derived NPCs as compared to the wild type cells. While several neuronal genes regulated by REST elevated in Ehmt1^-/+^ cells, expression of NRSF/REST mRNA was decreased. Mean fold change over wild type NPCs, n ≥ 3 independent experiments. (B-F) Western blot analysis of REST and H3K9me2 protein expression in (B) mECS following treatment with various concentrations of UNC0638 for 48h, (C) KS patient 1 iPSCs in comparison to control hiPSCs, (D) KS patient2 iPSCs in comparison to control hiPSCs, (E) EHMT1^-/+^ iPSCs in comparison to the isogenic control hiPSCs, (F) hiPSCs following treatment with various concentrations of UNC0638 for 72h, Quantification of Western blot analysis was performed by normalization to GAPDH, n ≥ 3. The expression of NRSF/REST protein was significantly decreased when the level/activity of EHMT1 was suppressed in all examined models. (G) qRT-PCR analysis of hiPSCs following treatment with 250nM UNC0636 for 72h to examine changes in the expression of REST-target genes; NRXN3, ACTA1 and Calbindin. Expression of REST-target genes were significantly elevated in the presence of UNC0636. Fold change over control untreated hiPSCs, n ≥ 3 independent experiments. Error bars represent s.e.m. \**P* < 0.05, \*\**P* < 0.01, \*\*\**P* < 0.001.

To establish whether this is a phenomenon common to KS patients, we generated hiPSCs from two patients (Figure S1). Both patient lines showed an approx. 2-fold decrease in H3K9m2, accompanied with an equivalent decrease of NRSF/REST protein (Figures 1C and D). As the microdeletion in KS patients ablates multiple genes, we employed CRISPR-Cas9 to introduce a 56-bp deletion in exon 12 of *EHMT1* gene to generate a hemizygous *EHMT1* knockout hiPSC line (*EHMT1*^-/+^), which would be isogenic with the parental wild type control cells. This deletion caused a loss of 50% of EHMT1 protein (Figure S2) and an accompanying reduction of H3K9me2 expression (Figure 1E). As seen in KS patient iPSC and mESC, reduced EHMT1 activity again led to a significant reduction in the expression of NRSF/REST protein in *EHMT1*^-/+^ cells (Figure 1E). Finally, as seen for wild type mESC, UNC0638 treatment of wild type hiPSCs caused a dose-dependent reduction of NRSF/REST protein, with half-maximal change at 250 nM (Figure 1F).

To confirm that changes to NRSF/REST protein lead to altered gene transcription, we examined expression of the NRSF/REST target genes NRXN3, ACTA1 and Calbindin(Bruce et al., 2004; Ballas et al., 2005; Wu and Xie, 2006), which normally commence expression in early stages of neurodevelopment. Using wild type human iPSC at the pluripotent cell stage, we found that 72 hours of UNC0638 treatment lead approximately 5-fold increase in mRNA levels of the NRSF/REST targets NRXN3, ACTA1 and Calbindin compared to treated cells (Figure 1G). Collectively, these mouse and human data indicate that EHMT1 regulates the level of NRSF/REST of pluripotent stem cells, and when reduced directly elevates expression of its downstream target genes.

### EHMT1 regulates NRSF/REST via suppression of miRNA-associated with psychiatric disorders

Conventionally, H3K9me2 is considered to be a transcriptional repressor (Ea et al., 2012), yet we observed that reduced EHMT1 activity leads to decreased NRSF/REST expression. To examine the possibility for a direct, positive effect of EHMT1 activity on *NRSF/REST* gene expression, we used ChIP-qRT-PCR analysis with a H3K9me2 antibody to search for evidence of EHMT1-mediated H3K9me2 marks. We found no evidence for H3K9me2 at the *NRSF/REST* promoter (Figures 2A and S3), arguing against an activating interaction of EHMT1 on *NRSF/REST* gene transcription.

We therefore considered a potential mechanism of de-repression of an intermediate NRSF/REST repressor. MicroRNAs (miRNAs) are ~22-nt noncoding RNAs expressed in a wide range of eukaryotic organisms (Bartel, 2009) and play a critical role in regulation of gene expression at the post-transcriptional level. The effects of miRNAs are mediated by post-transcriptional interactions with the 3’ untranslated region (3’UTR) of the target mRNA to induce mRNA decay or translational repression (Bartel, 2009). miRNAs play crucial roles at key stages in the development of the nervous system of various organisms. While a number of miRNAs can promote differentiation of neuronal stem cells and neural progenitors into specific neuronal cell types, eg. miR-9, others have been shown to induce proliferation of neuronal stem cells and neural progenitor, eg. miR-137 (Meza-Sosa et al., 2014). Interaction between cell type specific transcription factors and miRNAs are required for neural stem cells to differentiate into functionally mature neurons. Brain-related miRNAs including miR-142, miR-153 and miR-9 and have been shown to target NRSF/REST mRNA (Wu and Xie, 2006). and hence offer a potential pathway connecting repressive EHMT1 histone methylation and NRSF/REST regulated gene expression.

To search for miRNAs regulated by EHMT1, we performed miRNA-seq to identify the miRNA species that are differentially expressed when EHMT1 activity is inhibited with UNC0638. We detected 56 miRNAs with greater than 2.5-fold increase of expression in the presence of EHMT1 inhibitor (Figure S4). 11 of these miRNAs were predicted to target NRSF/REST mRNA based on miRDB database, and 9 replicated by qRT-PCR analysis of UNC0638-treated hiPSC, including miR-142, miR-153-1, miR-26a-2, miR-548f-1 (upregulated by 20.6 ± 3.50, 6.70 ± 0.19, 7.95 ± 1.908 and 4.62 ± 0.35-fold, respectively) (Figure 2B). We further examined whether the association between EHMT1 activity and miRNA expression was conserved in mESC and observed expression of miR-142, miR-153-1, miR-181a and miR-769 were all elevated after UNC0638 treatment (Figure 2C). To investigate direct linkage between these NRSF/REST-regulating miRNAs and EHMT1 activity we examined their association with H3K9me2-modfied chromatin. We selected three miRNAs, miR-142, miR-153-1 and miR-26a-2, whose expression was significantly upregulated when EHMT1 is inhibited, designed PCR primers to their transcription start sites (TSSs) and non-TSSs sites as negative controls (Figure 2A). H3K9me2-ChIP in in hiPSC showed that the TSS of all three miRNA were modified (Figure 2A). Our combined gene expression and ChIP analysis indicates indirect regulation between EHMT1 and NRSF/REST, acting via suppression of intermediate miRNA that mediate NRSF/REST mRNA

**Figure 2:**
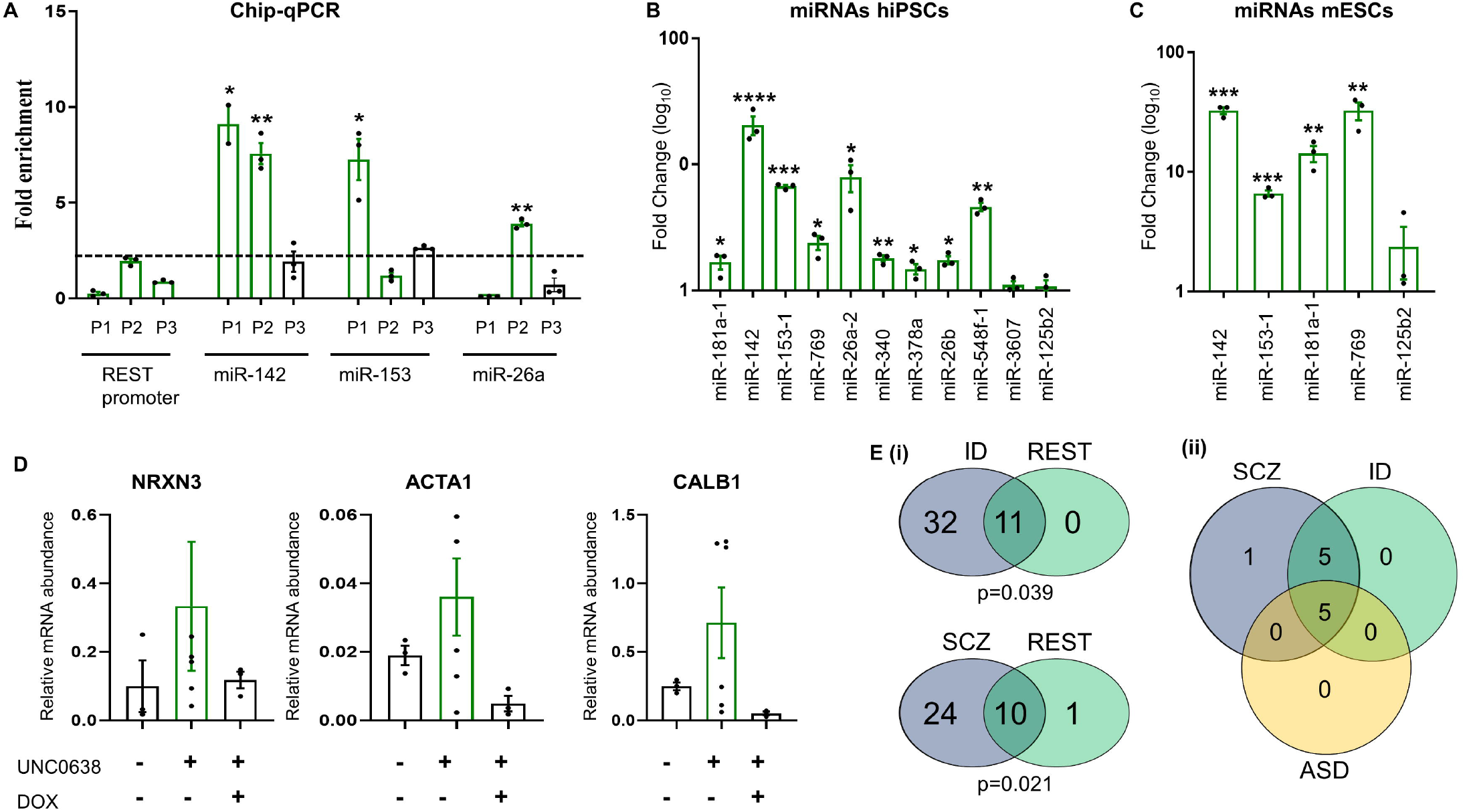
EHMT1 suppresses expression of miRNA. (A) ChIP-seq analysis of NRSF/REST and miRNA genes. ChIP-qRT-PCR analysis of H3K9me2 modification within 3 regions around NRSF/REST promoter, (P1, P2 and P3) and around (P1 and P2) and away (P3) from the TSSs of miR-142, miR-153 and miR-26a (see Fig S3). ChIP was performed with an anti-H3K9me2 antibody, and H3K9me2 enrichments were analysed by qRT-PCR. Enrichments (relative to the input DNA in specific genomic regions) were assessed by ChIP. The primer sets are listed in Supplementary Table S4. H3k9me2 was accumulated at miR-142 miR-153 and miR-26a TSS sites, whilst regions around the NRSF/REST promoter were not occupied by H3k9me2. (B,C) miRNA-seq validation by qRT-PCR. To confirm the obtained miRNA-seq results, qRT-PCR analysis was performed to validate the differentially upregulated miRNAs in UNC0638-treated (B) hiPSCs and (C) mECS. The qRT-PCR data mirrored those of the miRNA-seq data, n ≥ 3. Mean±SEM \**P* < 0.05, \*\**P* < 0.01, \*\*\**P* < 0.001, \*\*\*\**P* < 0.0001. (D) Relative mRNA abundance measured by qRT-PCR of the REST-target genes NRXN3, ACTA1 and Calbindin1 (CALB1) in wild type hiPSCs treated with UNC0638. Induction of RESTDUTR expression with doxycycline (DOX) suppresses the up-regulation of NRXN3, ACTA1 and Calbindin seen in the presence of UNC0638. (E) (i) Venn diagrams showing association between miRNAs following UNC0638 treatment and those associated with Intellectual Disability (ID) and schizophrenia (SCZ). Crossover probability was assessed for each disorder by calculating hypergeometric probability, with P□<□0.05 considered statistically significant (ii) Venn diagram to show overlap of REST targeting miRNAs between ID, SCZ and ASD.

To test our mechanistic hypothesis, we introduced a doxycycline-inducible (“tet-on”) expression plasmid into the AAVS1 safe harbour site of wild type hiPSC to express a recombinant NRSF/REST gene (RESTDUTR) that lacks the miRNA-target region within the 3’UTR region of its mRNA (Figure S5). The rationale for this strategy is that removing all miRNA target sites in the NRSF/REST mRNA would overcome the redundancy due to the presence of multiple EHMT1-regulated miRNAs. For pluripotent cells in the presence of UNC0638, but absence of doxycycline, we observed an elevation of NRXN3, ACTA1 and Calbindin gene expression as previously seen for wild type hiPSCs (Figure 2E). Expression of RESTDUTR following addition of doxycycline blocked the UNC0638-induced elevation of NRXN3, ACTA1 and Calbindin gene expression (Figure 2E). This result demonstrates that uncoupling of NRSF/REST expression from its miRNA regulators is sufficient to overcome the effects of reduced EHMT1 activity.

Given the potential association between EHMT1 activity and patient diagnosis that extend across a range of NDD, we examined the relationship between the miRNA up-regulated due to EHMT1 inhibition and GWAS for ID, schizophrenia and ASD. Remarkably, 43 and 34 of the 56 up-regulated miRNA have genetic-association with ID and schizophrenia respectively (Table S5). In contrast, only 15 ASD-associated were up-related, all of which overlapped with the ID gene set and 13 of which were also associated with schizophrenia (Figure S6). Of the up-regulated miRNA gene sets that are NRSF/REST regulators, there was significant enrichment for those associated with ID (all 11 miRNA) and schizophrenia (10 of 11 miRNA), but not ASD ((5 of 11 miRNA) (Figure 2E, S6). Furthermore the five NRSF/REST targeting miRNA associated with ASD where present in both ID and schizophrenia overlaps (Figure 4E), giving a core miRNA gene set of miR-26a, miR-26b, miR-153, miR-181a and miR-548. This analysis suggests a broad association of this miRNA-mediated NRSF/REST regulatory pathway and psychiatric risk, particularly for a diagnosis of ID and schizophrenia.

### Reduced EHMT1 activity accelerates neuronal differentiation

We investigated whether the relationship between reduced EHMT1 activity and NRSF/REST protein is maintained beyond the pluripotent cell state and persists into neurodevelopment. Mutant *EHMT1*^+/-^ hiPSCs and their isogenic wild type controls were differentiated into neurons using a standard dual-SMAD inhibition protocol (Chambers et al., 2009) and NRSF/REST protein levels sampled at time points that span neuronal differentiation (Figure 3A). NRSF/REST protein was significantly lower at several time points of differentiation in *EHMT1*^-/+^ mutant cells compared to isogenic controls. This indicates that the effect of EHMT1 on NRSF/REST protein persists from early NPC stage into neuronal differentiation. The reduction of NRSF/REST in differentiating *EHMT1*^-/+^ iPSC was accompanied by increased expression of the human orthologues of the target genes MASH1 and NGN2, which are first expressed in NPC (Ballas et al., 2005; Parras et al., 2002) (Figures 3B and C). Expression of MASH1 in control hiPSCs derived NPCs was also increased in the presence of UNC0638 and suppressed by doxycycline-mediated induction of RESTDUTR mRNA (Figure 3D).

**Figure 3.**
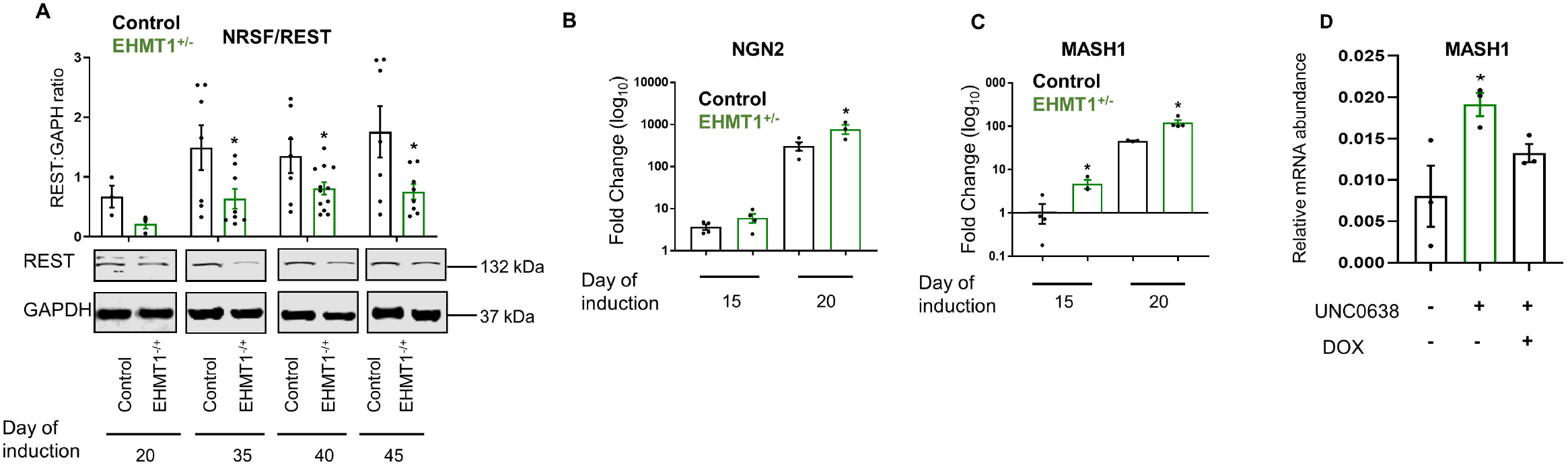
Reduced EHMT1 activity elevates REST-target gene expression during neuronal development. (A) Western blot analysis of NRSF/REST protein expression in hiPSCs-derived neurons at Day 20 (NPC stage) and maturing neurons at Days 35, 40 and 45 of differentiation. NRSF/REST expression decreased in EHMT1^-/+^-derived neurons. Quantification of Western blot analysis was performed by normalization to GAPDH. A representative image from at least three independent experiments is shown. (B,C) Time-course qRT-PCR analysis at Days 15 and 20 of differentiation to examine changes in the expression of lineage-specific REST-target genes, (B) NGN2 and (C) MASH1 in EHMT1^-/+^-derived neurons in comparison to their expression in the isogenic control hiPSCs. Expression of NGN2 and MASH1 increased in EHMT1^-/+^-derived neurons. Fold change over day 0 of differentiation, n ≥ 3 independent experiments. (D) Relative mRNA abundance measured by qRT-PCR of MASH1 in wild type hiPSCs treated with UNC0638. Induction of RESTDUTR expression with doxycycline (DOX) suppresses the up-regulation of MASH1 seen in the presence of UNC0638.

Our results suggest that EHMT1 acts via NRSF/REST as a negative regulator of neuronal gene expression to prevent premature neurodevelopment. We therefore investigated whether this impacted on the differentiation of neurons in general and hence not restricted to the subset of NRSF/REST-regulated genes. As seen for known direct NRSF/REST target genes, mRNA of the non-NSRF/REST target genes, Pax6, NCAM and Nestin (Sansom et al., 2009)’(Suzuki et al., 2010) were also elevated at Day 10 and 20 of neuronal differentiation as cells transition from the NPC stage (Figures 4A, B and C), suggesting a general acceleration of neuronal development. Wild type hiPSCs differentiated into neurons in the presence of UNC0638 again had increased expression of the non-NRSF/REST target neuron genes Nestin, an increase that was suppressed by doxycycline-induced RESTDUTR expression (Figure 4D). These results were indicative of accelerated neuronal development. To pursue this further, we examined MAP2 (microtubule-associated protein 2), protein that is expressed in neurons (Harada et al., 2002) during neuronal differentiation. MAP2 mRNA was elevated in *EHMT1*^+/-^ cells during NPC stages indicative of a rapid transition through the progenitor cell state and into full neuronal differentiation (Figure 4E). As also seen with other genes, MAP2 expression was increased in wild type hiPSCs differentiated in the presence of UNC0638 (Figure 4F) and again this was suppressed by doxycycline-induced RESTDUTR expression (Figure 4G). These results indicate accelerated neuronal differentiation in *EHMT1*^-/+^ hiPSCs, commencing as cells leave the pluripotent cell state and continues to the formation of mature neurons. This is consistent with similar observations of *in vitro* neurodevelopment of NRSF/REST hemizygous mESC mutations (Aoki et al., 2012).

**Figure 4:**
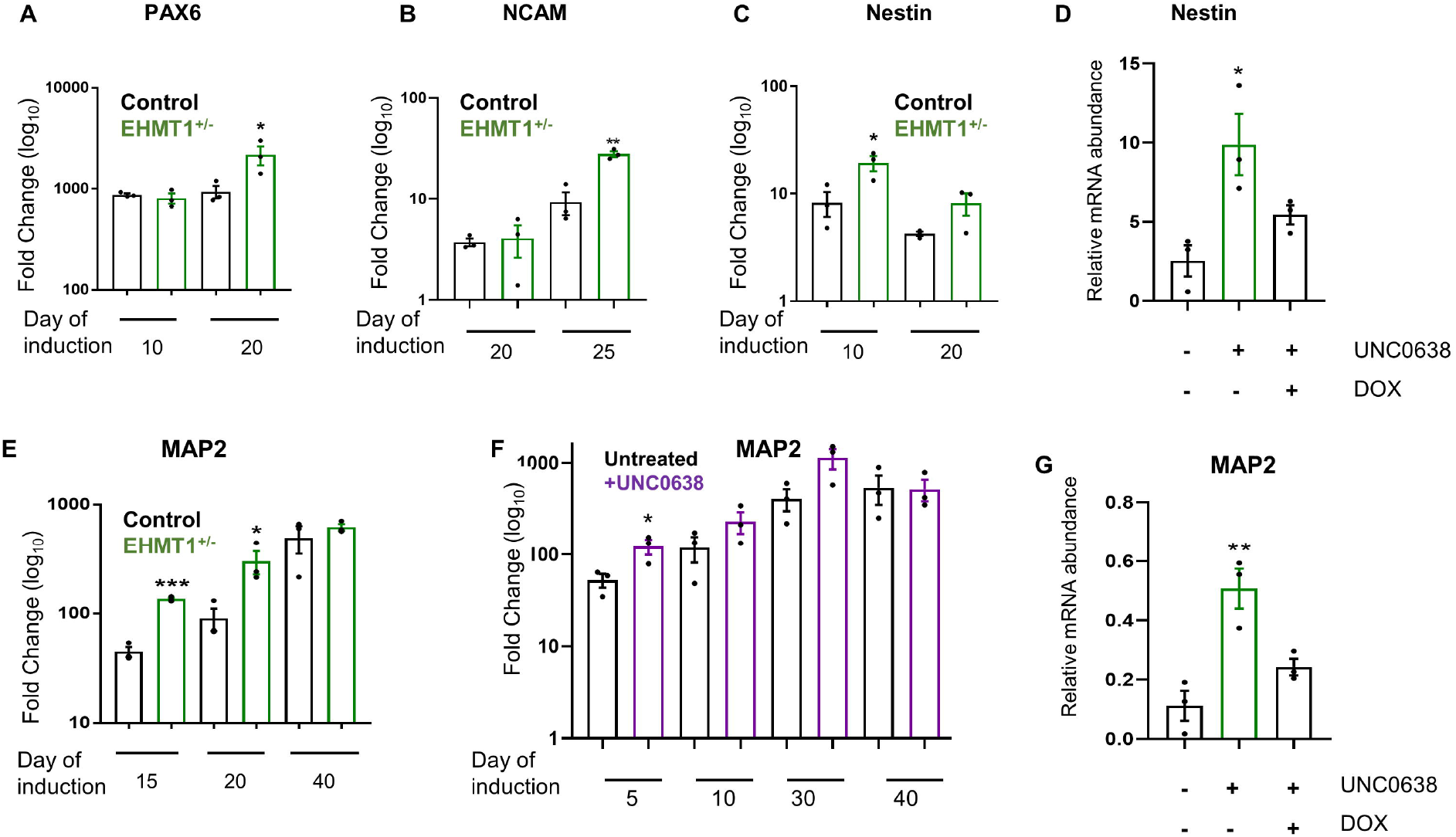
Reduced EHMT1 activity results in accelerated neuronal development. (A-C) Time-course qRT-PCR analysis at days 10 and 20 of differentiation to examine the expression of the neural progenitor markers, (A) PAX6, (B) NCAM and (C) Nestin in EHMT1^-/+^-derived neurons relative to their expression in the isogenic control NPCs. (D) Relative mRNA abundance measured by qRT-PCR of Nestin in wild type hiPSCs treated with UNC0638. Induction of RESTDUTR expression with doxycycline (DOX) suppresses the up-regulation of Nestin seen in the presence of UNC0638. (E) Time-course qRT-PCR analysis at days 15, 20 and 40 of differentiation to examine changes in the expression of the neuronal marker MAP2 in EHMT1^-/+^-derived neurons relative to their expression in the isogenic control neurons. Fold change over day 0 of differentiation, n ≥ 3 independent experiments. Error bars represent mean ±SEM. \**P* < 0.05, \*\**P* < 0.01, \*\*\**P* < 0.001. (F) Time-course qRT-PCR analysis at days 5, 10, 30 and 40 of differentiation to examine the change in the expression of the neuronal marker, MAP2 in hiPSCs-derived neurons treated with UNC0638 (250nM). The expression of MAP2 increased in UNC0638-treated neurons as compared to untreated neurons. (G) Relative mRNA abundance measured by qRT-PCR of MASH1 in wild type hiPSCs treated with UNC0638. Induction of RESTDUTR expression with doxycycline (DOX) suppresses the up-regulation of MAP2 seen in the presence of UNC0638.

### Rapid neurodevelopment as a consequence of EHMT1 inhibition is associated with increased apoptosis

Although we observed a consistent pattern of rapid neurodevelopment in cells with reduced EHMT1 activity in early developmental stages by Day 40 these differences appeared to be lost (Figures 4E, F and S7). Previous reports described mice lacking NRSF/REST as having a transient increase in neurogenesis but eventually leading to decreased granule neurons production (Gao et al., 2011). We therefore investigated the impact of the prolonged inhibition of EHMT1 activity on the neuronal differentiation.

We monitored neuronal cell number using cell staining with the neuronal nuclear marker, NeuN. Wild type cells were differentiated to neurons in the presence and absence of UNC0638 and sampled at Days 25, 30, 35 and 45 (Figures 5A-C). Comparison of the number of NeuN-stained cells treated with UNC0638 and untreated controls demonstrated a progressive decrease in the percentage of NeuN stained cells in the presence of the inhibitor. By Day 45, UNC0638-treated cell cultures had 50% of the number of NeuN positive cells compared to untreated controls (Figure 5C). This was consistent with the levelling of MAP2 expression between control and cells with reduced EHMT1 at later developmental time points (Figures 4E and F).

**Figure 5:**
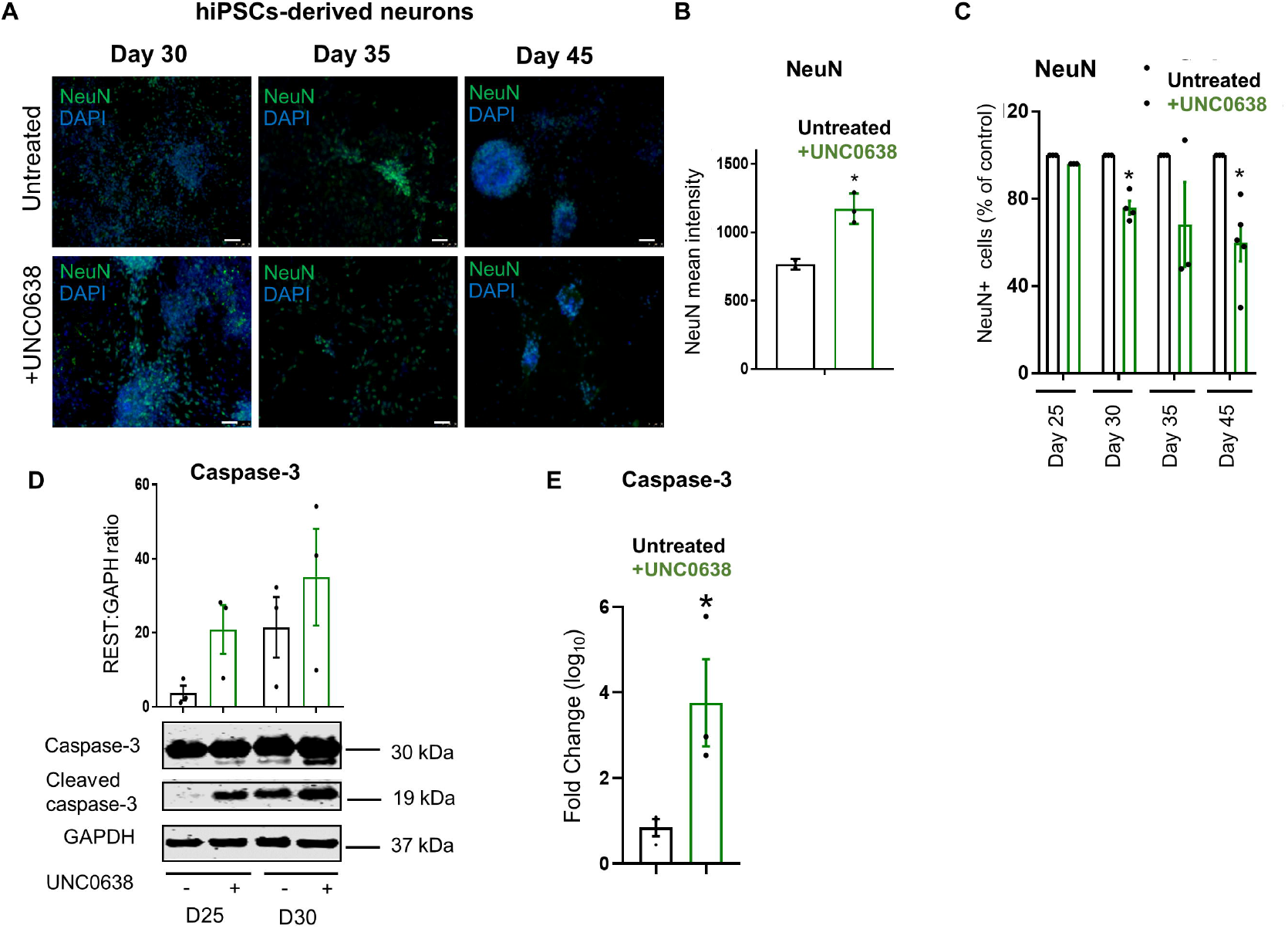
Adjustment of neuronal cell numbers during later differentiation. (A) Representative immunocytochemistry images of hiPSCs-derived neurons at days 30, 35 and 45 of differentiation stained with NeuN (green) and counterstained with DAPI (blue). Scale Bar, 50μM. (B) NeuN expression (mean average intensity of cell nuclei) of hiPSCs-derived neurons treated with UNC0638 (250nM) at day 35 of differentiation. Mean average intensity of NeuN protein was significantly increased in the presence of UNC0638. (C) Total number of cells stained with NeuN in hiPSCs-derived neurons at days 25, 30, 35 and 45 of differentiation in the absence and presence of UNC0638. The number of cells stained with NeuN was significantly reduced in the presence of UNC0638 after Day 30, n ≥ 3 independent experiments. (D) Western blot analysis at days 25 and 30 to examine the expression of cleaved-caspase-3 in the presence and absence of UNC0638 (250nM). Top panel represents the expression of total caspase-3, middle panel represents the expression of cleaved-caspase-3 and bottom panel represents the expression of GAPDH. Quantification of Western blot analysis was performed by normalization to GAPDH. Expression of cleasved-caspase-3 elevated in the presence of UNC0638, n ≥ 3. (E) qRT-PCR analysis to examine the change in the expression of caspase-3 in hiPSCs-derived neurons treated with UNC0638 (250nM). The expression of caspase-3 increased in UNC0638-treated neurons as compared to untreated neurons. Fold change over untreated neurons (mean), n ≥ 3 independent experiments. Error bars represent SEM. \**P* < 0.05.

During neurodevelopment, disturbance of neuronal activity can potentially lead to deregulation of neuronal survival (Dekkers et al., 2013). To determine the mechanism underlying the neuronal loss, we examined Caspase-3 activation in UNC0638-treated cells in relation to the control. At days 25 and 30 of neuronal differentiation, which are the time points preceding the reduction in cell number, we detected an elevated Caspase-3 cleavage to the activated form in UNC0638-treated cells (Figure 5D). The increase in Caspase-3 activation was accompanied by increased Caspase-3 gene expression in control neurons treated with UNC0638 (Figure 5E). These observations support the hypothesis that in *EHMT1*^-/+^ cells the reduction in the number of cells stained with NeuN and the neuronal gene expression may be due to an induction of programmed cell death.

### Reduced EHMT activity leads to aberrant neuronal function

We next examined whether the abnormal developmental programme seen in *EHMT1*^+/-^ mutant cells altered neuronal function. Functional maturation of hiPSCs-derived neurons has been previously described (Shi et al., 2012b), with firing action potential observed at approximately 50-100 days of differentiation (Shi et al., 2012b). To investigate whether shortterm inhibition of EHMT1 activity leads to abnormal neuronal functionality, neuronal activity was detected following inhibiting EHMT1 activity for ~two weeks. hiPSCs were transformed into neurons using our standard protocol and when the neurons begin to form ~day 35, N2B27 medium was switched to BrainPhys medium, as this medium supports the neurophysiological properties of the neurons (Bardy et al., 2015). At this stage, cells were either treated with UNC0638 (250nM) to inhibit EHMT1 activity or maintained as untreated controls. Spontaneous calcium influx was measured two weeks later when the neurons typically begin to exhibit firing action potential and the cultures become more physiologically active (Shi et al., 2012a). The number of spontaneously active cells in UNC0638-treated cultures was significantly elevated compared to untreated controls (Figure 6A). Furthermore, the frequency of calcium events per neuron was significantly higher in cells treated with UNC0638 compared to the untreated culture (Figures 6A and B). This indicates that the neurons derived from EHMT1 inhibited cells exhibit accelerated maturation and increased excitability in relation to the control cells, which recapitulates the finding of the elevated levels of neuronal markers observed in *EHMT1*^-/+^ cells.

**Figure 6:**
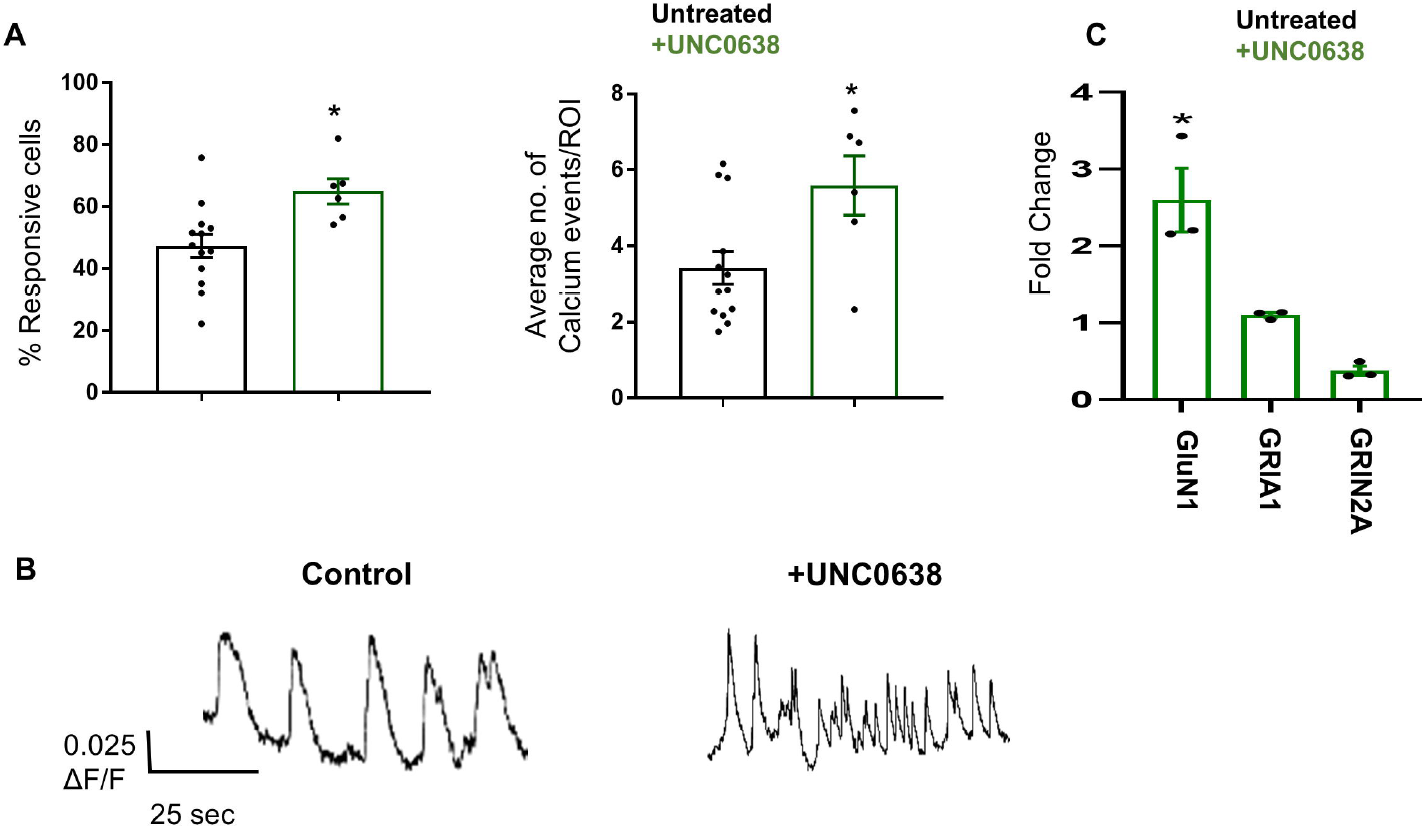
Reduced EHMT activity leads to aberrant neuronal activity. Calcium imaging of hiPSCs-derived neurons. Differentiated neurons were analysed for spontaneous calcium events using Cal-520tm AM. Neurons were exposed to the dye and they were imaged for fluorescence with a 488 nm excitation filter. Images were captured at 10 Hz and recorded for 5 minutes. (A) The percentage of responsive cells and average calcium events/ROIs for control untreated hiPSCs and cells treated with UNC0638. Inhibiting EHMT1 activity increased the percentage of responsive cells as well as the average calcium event frequencies of hiPSCs-derived neurons. (B) Representative traces of calcium influx of neuronal cultures in the presence and absence of UNC0638 (250nM). Neurons were treated with UNC0638 for two weeks and calcium imaging was performed around day 50 of differentiation to compare the number of calcium event frequencies between control neurons and those treated with UNC0638. Vertical scale bar in f shows 0.025 (ΔF/F); horizontal bar shows 25s. N = 300 ROIs for all cell lines across 2 imaging regions over 3 coverslips/line. (C) qRT-PCR analysis to examine changes in the expression of GluN1, GRIA1 and GRIN2A in EHMT1^-/+^-derived neurons relative to their expression in isogenic control neurons. The expression of GRIN1 was significantly elevated in EHMT1^-/+^ cells, while those of GRIA1 and GRIN2A were unchanged, n ≥ 3. Error bars represent SEM, \**P* < 0.05.

Frega and coworkers (2019) described aberrant network activity of neurons derived from KS patient iPSCs in comparison to control neurons, that is driven by NMDA receptor (NMDAR) subunit 1 hyperactivity. The authors detected significant elevation in the level of the gene encoding NMDAR subunit 1 *(GluN1)*. We therefore investigated changes in the expression of GluN1, the other NMDAR subunit *GluN2A* and an AMPAR subunit *GRIA1*. We observed a significant upregulation in the expression of GluN1 mRNA in EHMT1^-/+^ iPSCs-derived neurons. These results are consistent with those reported by Frega and coworkers (Figure 5C). Elevated GluN1 may lead to aberrant NMDA-mediated glutamate signalling. Expression of GRIA1 did not increase and in contrast to our observations in mESC (Figures 1A and 5C) we observed little change for GluN2A.

## Discussion

Here, we report a biological pathway that connects the molecular lesion in EHMT1 activity to altered neuronal cell development and neuronal function in human iPSC-derived neurons. The key mechanistic components of the pathway are the transcriptional regulator NRSF/REST and its control via miRNA, with reduced EHMT1-mediated H3K9me2 resulting in elevated miRNA transcription. As a consequence, gene expression of both NRSF/REST-regulated and general neuronal specific markers is elevated with lower levels of EHMT1 activity, but can be reduced to control levels by expression of a NRSF/REST mRNA that lacks the miRNA regulation sites. Although KS presents a strongly genetically penetrant case for this pathway, our genetic association analysis suggests a broader association for ID and schizophrenia, with implications for therapeutic intervention.

NRSF/REST plays a key role in repression of neuronal gene expression to maintain stem cells in the undifferentiated state (Schoenherr and Anderson, 1995) (Ballas et al., 2005), making it a major regulator of neurogenesis and neural differentiation (Yang et al., 2012). Here, we can explain the *in vitro* cell phenotype of KS patient cells as due to down-regulation of NRSF/REST. Our observations fit with previous investigation of NRSF/REST hypofunction in neurodevelopment (Schoenherr and Anderson, 1995). In the mouse brain, conditional NRSF/REST knockout mice show rapid neuronal differentiation of hippocampal neural stem cells and elevation in the expression of pro-neuronal genes, *NeuroD1, Tuj1, and DCX* (Gao et al., 2011). In the study reported here, we show that reduced NRSF/REST expression in EHMT1^-/+^ cells is associated with elevated expression of the human pro-neural transcription factors MASH1 and NGN2.

In human neuronal culture, decreased nuclear NRSF/REST has been observed in neuronal cultures derived from sporadic Alzheimer’s Disease (AD) patient cells and again leads to accelerated neural differentiation and increased excitability, which can be reversed by exogenous NRSF/REST expression (Meyer et al., 2019). In the context of NDD, Down’s Syndrome cells have increased expression of DYRK1A which leads to reduced NRSF/REST and mis-regulation of neurodevelopmental genes (Lepagnol-Bestel et al., 2009). Likewise, suppression of Chromodomain helicase DNA-binding protein 2 (CHD2), associated with a range of NDD, including ASD and ID, was shown to inhibit the self-renewal of radial glial cells and increase the generation of neural progenitors and neurons and this phenotype was attributed to the reduction occurred in the expression of the neuronal regulator NRSF/REST (Shen et al., 2015).

In our EHMT1^+/-^ hiPSC cultures, in addition to an accelerated neurodevelopmental programme, we observed a loss of neuronal cells due to elevated apoptosis as neuronal maturation began. The exact mechanism leading to cell death at these later stages is unclear and may be a consequence of *in vitro* culture having an increased sensitivity to apoptotic signals compared to the brain environment, but it is noteworthy that NRSF/REST-target genes include cell death inducing genes and may directly induce apoptosis (Lu et al., 2014). We also note that in the mouse brain, both loss of *ehmt1* (Benevento et al., 2017) or *rest* genes (Bibel et al., 2004b) increase cell proliferation and adult neurogenesis, and that prolonged loss of NRSF/REST leads to a functional depletion of the adult neuronal stem cells and decreased granule neurons production (Gao et al., 2011). Furthermore, analysis of post-mortem AD, where NRSF/REST is reduced compared to age matched controls, show elevation of NRSF/REST targets, including genes encoding pro-apoptotic signalling components, associated with neurodegeneration (Lu et al., 2014).

Our results also show that the NRSF/REST gene is not a direct target of EHMT1, but instead is regulated via control of miRNA expression. This places NRSF/REST as a key node within a miRNA-mediated gene regulatory network. Previous reports described a large repertoire of miRNAs that regulate NRSF/REST mRNA stability (Gao et al., 2012; Duan et al., 2014). There is a considerable enrichment for miRNA that target and degrade NRSF/REST mRNA almost those upregulated in hiPSCs in the pluripotent state. The high degree of multiplicity in the number of miRNAs targeting NRSF/REST makes this regulation robust and it is unlikely that targeting a single miRNA could be sufficient to reverse the effects of KS. Therapeutic intervention would need to be directed at upstream of miRNA transcription, perhaps by stabilising H3K9me2-mediated regulation; destabilising the miRNA target sites in the 3’UTR or reducing NSRF/REST protein turnover. Some miRNAs are both regulators of NRSF/REST and also regulated by NRSF/REST, creating a potential feedback loop. This is the case for miR-26a and miR-181a in the subset of EHMT1 sensitive, pluripotent miR genes, and may act to maintain NRSF/REST mRNA at a constant level. In contrast, miR-142 and miR-153 are not known to be repressed by NRSF/REST and therefore have potential to drive NRSF/REST expression down, pushing it into a state of low-level expression in response to reduced H3K9me2. Finally, miRNA have multiple gene targets, diversifying the EHMT1 signal throughout the gene regulatory network, and raising the possibility that other cell signalling components may be regulated in parallel to NRSF/REST.

The NRSF/REST gene or its regulatory miRNAs have not to date been genetically associated with psychiatric disorders. This may be because of a combination of network robustness protecting against minor deficits in its upstream regulatory pathway, and the severity caused by major changes in NRSF/REST expression, as reported with its association with dementias (AD, Huntington’s Disease and Parkinson’s Disease), ischemic shock and some NDD. This is not however the case for its regulatory cofactors and downstream targets. NRSF/REST exerts its epigenetic repression via engaging a number of regulatory proteins with their chromatin targets. In particular KMT1C (EHMT2), which associates with NRSF/REST, has co-morbidity between ID and ASD (Koemans et al., 2017). Further NRSF/REST-associated genes have also been shown to link to ID and ASD, as have a range of other epigenetic regulators. Our cross-disorder analysis of the miRNA under EHMT1-regulation and their association with GWAS-significant miRNA genes suggests hitherto unexplored linkage with NSRF/REST not only for KS but across a broader range of ID and schizophrenia cases. The consideration of NRSF/REST regulation within the biological pathways leading to NDD may offer significant insights for therapeutic strategy.

In summary, this study identifies a mechanism linking EHMT1 activity to the neuronal regulator NRSF/REST through miRNA-dependent pathway that leads to altered neurodevelopment. It offers a mechanism for the specific case of KS but also reveals the presence of a more extensive pathway centred around NRSF/REST regulation of the neurodevelopmental gene regulation programme.

## Method details

### Human iPSCs culture and neuronal differentiation

Human (h)iPS cells were maintained on matrigel (Corning) using Essential 8™ medium (ThermoFisher scientific) at 37°C, 5% CO2 in an incubator (Galaxy 170R, New Brunswick, USA) (Nishishita et al., 2015). Medium was changed every day and cells were passaged using gentle cell dissociation reagent (Stemcell Technologies) or singularised using accutase (Stem Cell Technologies). To promote cell survival during enzymatic passaging, cells were passaged with the p160-Rho-associated coiled-coil kinase (ROCK) inhibitor Y-27632 (10 μM; Tocris) or RevitaCell™ Supplement (ThermoFisher scientific) (Watanabe et al., 2007).

Cell morphology was continually analysed and cells were discarded if the morphological characteristics of stem cell colonies were not maintained or if staining for routine pluripotency markers was negative (Figure S1A). In addition, cells were regularly tested for mycoplasma using the MycoAlert™ plus mycoplasma detection kit (Lonza).

hiPSCs were differentiated into glutamatergic neurons based on the method reported by Chambers et al with some modifications (Chambers et al., 2009). Briefly, prior to differentiation, cells were passaged onto 12-well plates coated with growth factor-reduced Matrigel (Corning). Once cells were >80% confluent (D0), medium was switched to N2B27 (2/3 DMEM/F12; 1/3; neurobasal; B27-RA; N2; 1xPSG; 0.1mM β-mecaptoethanol) + 100nM SB431542 and 100nM LDN193189 and half of the medium was changed every other day. After 8 days cells were passaged onto fibronectin coated 12-well plates at a ratio of 2:3 in un-supplemented N2B27 media. After a further 10 days, cells were passaged onto PDL (Sigma)/laminin (Roch) coated 12-well plates as single cells at a density of 200000 cells/cm^2^. Media were switched to N2B27 with B27+ retinoic acid in place of B27-RA, and 10μM DAPT for 7 days, followed by using N2B27 with B27 + retinoic acid for the remainder of the culture.

### Kleefstra syndrome (KS) hiPSCs generation

iPSCs derived were derived from two Kleefstra syndrome (KS) patients: a 22-year-old female patient, non-verbal (IQ not measurable), with a history of epilepsy (treated with carbamazepine and levetiracetam), hypotonia, anxiety disorder and depression; and a 20 female patient 20, (FSIQ 53) diagnosed with ASD, specific phobia, psychotic symptoms, hypotonia, unprovoked seizures; and cardiac, mitral valve insufficiency. The participants were recruited as part of a research cohort on neurodevelopmental copy number variants at Cardiff University (the Defining Endophenotypes from Integrated Neuroscience [DEFINE] Study). Procedures included clinical and cognitive testing, where possible, and blood sampling for generation of iPSCs and were approved by the South East Wales Research Ethics Committee. Where participants did not have capacity to consent, as in this case, a representative (next of kin) provided written informed consent on their behalf. Peripheral blood mononuclear cells (PBMCs) from the donor were isolated, expanded, and reprogrammed using CytoTune-iPS 2.0 Sendai reprogramming kit (A16517, ThermoFisher scientific) following the manufacturer’s instructions, and as previously described (Chichagova et al., 2016). The IBJ4 cell line we used as the control line derived from the BJ fibroblast cell line (ATCC; CRL-2522), when not indicated otherwise.

KS iPSCs karyotyping was performed by treating the cells with Demecolcine solution (Sigma) and then processed with standard methods. Chromosomes were stained with DAPI and the images obtained by a Leica DMI6000b fluorescent microscope in 100x objective. Karyotype analysis was performed and a minimum of 20 metaphases were evaluated. Analysis showed 46, XX normal diploid female karyotype (Figure S1B). The chromosomes were classified according to the International System for Human Cytogenetic Nomenclature (ISCN), and each patient cell line possessed a 9q34 deletion.

### Mouse ES cells culture and neuronal differentiation

Mouse ES cells (mESCs) were purchased from EUCOMM and the European Mouse Mutant Cell Repository Centre (EuMMCR). Two clonal populations of Ehmt1 mutant cells were used. Cells had a single copy of the Ehmt1^tm1a(EUcOMM)Hmgu^ allele, which is a ‘knockout first’ conditional allele (Skarnes et al., 2011). The wildtype control line was generated using the flp-allele to restore a wild type gene. mESCs were maintained on gelatin-coated plates in knockout DMEM (Gibco), supplemented with ESC certified FBS (Invitrogen), L-Glutamine (Gibco), 2-mercaptoethanol (Sigma) and ESGRO leukemia inhibitory factor (LIF) (Chemicon) at 37°C in an incubator (Galaxy 170R, New Brunswick, USA). The mECs media were changed daily and cells were passaged every other day using TrypLE (Gibco) as a dissociation agent. mESC were expanded on a population of mouse embryonic fibroblast cells (MEFs) for two or three passages, then on gelatin-coated plates for three or four further passages before plating without gelatin on non-adhesive bacteriological plates in media lacking LIF for 4 days, where cells form cellular aggregates. 5 μM Retinoic acid was added to the culture until day 8. At this stage, cells were dissociated with 0.05% trypsin (Sigma) and seeded at 1.5 x 10^5^ per cm^2^ density on Poly-D-ornithine (Sigma)/laminin (Roch) in N2 medium (Sigma) for 48h. Subsequently, N2 medium was switched to complete medium (Sigma) (Bibel et al., 2004a), and cells were maintained in this media for the remainder.

### Generation of Stable CRISPRn and REST iPSC Lines

iPSCs were dissociated with accutase, resuspended in phosphate buffered saline (PBS), and counted using a hemocytometer. For plasmid (CRISPRn/REST) transfections, 4D-Nucleofector™ unit X Kit (program CA-137; Lonza) was used according to the manufacturer’s instructions. To generate the Doxycycline (Dox)-inducible CRISPRn and inducible REST iPSC lines, one million iPSCs were nucleofected with appropriate vector (2.5 μg), carrying the puromycin resistance gene cassette and each AAVS1 TALEN pair (1 μg) (Mandegar et al., 2016). Cells were then seeded in six-well plates in Essential 8™ medium supplemented with Y-27632 (10 μM). Selection was applied 3 days post-nucleofection using 0.5 mg/ml puromycin (Life Technologies). Selection was maintained for 10 days until stable colonies appeared. Colonies with a diameter of greater than ~500 μm were manually picked using a P200 pipette tip and transferred to individual wells of a 24-well plate containing Essential 8™ medium supplemented with Y-27632 (10 μM). Clones were then expanded into larger vessel formats.

### Syntactic gRNA transfection

Syntactic gRNA transfection was performed as previously described (González et al., 2014). Briefly, iCas9 hiPSCs were treated with 2 μg/ml Dox one day before transfection. On the day of transfection, cells were transfected with Synthetic gRNA (SygRNA) (crRNA and transactivating crRNA (tracrRNA) guide) using Lipofectamine CRISPRMAX (Life Technologies) following the manufacturer’s instructions. 10 pmols gRNA were used and both SygRNA and Lipofectamine CRISPRMAX were diluted separately in Opti-MEM (Life Technologies), before being mixed together, incubated for 5 min at room temperature (RT), and added dropwise to cultured hiPSCs. A second transfection was performed, in a similar manner 24 hr later, to increase the editing efficiency. To test for successful editing, genomic DNA was extracted using QuickExtract ™ DNA Extraction Solution (Illumina, USA), and PCR amplification using primers encompassing the CRISPR target site, 5’-AGCAGCATCTCTCACCGTTT-3’ and 5’-CTTTTTCAGGTGGACGACTGG-3 was performed. PCR products of ~250-bp were size-separated by electrophoresis on a 4% agarose gel to check for any insertions or deletions which might be arisen as a result of CRISPR editing.

### CRISPR/Cas9 off-target detection

CRISPR/Cas9 off-target mutations were also considered. To that end, we used the open CRISPR design tools (Sigma) to predict the potential off-target sites which share up to 3 base mismatches with the sequence of our guide RNA. We have not identified any sites which have 1 or 2 base mismatches with our guide. However, we found four sites which have 3 base mismatches with the sequence of our guide. We have tested those regions using appropriate primers encompassing the potential edit sites, and a list of primers used and the product sizes are shown in Table S2. These off-target sites were amplified and tested for any insertions or deletions on a 4% agarose gel.

### Inducible REST vector design

To generate inducible custom REST vector, Dox-inducible CRISPR nuclease (CRISPRn) plasmid (pAAVS1-PDi-CRISPRn, Addgene) was used as a starting material. The Cas9 gene was digested with AfIII and AgeI restriction enzymes and the linearised vector, lacking Cas9 gene, was purified using gel clean-up kit (Wizard^®^ SV Gel and PCR Clean-Up, A9281, Promega) (Figure S4B). REST cDNA (~3300bp) missing the 3’URT (GenBank BC132859.1) was synthetised as a double-strand DNA using IDT service (https://www.idtdna.com/pages/products/genes/gblocks-gene-fragments). REST gene was <u>prepared as two fragments of ~1500bp with 40-bp overlaps at either end</u> which are homologous to the CRISPRn backbone vector (Figure S4C). The fragments were obtained as lyophilised DNAs, which was reconstituted following the manufacturer’s instructions.

NRSF/REST fragments as well as the linearised backbone vector were assembled in an isothermal Gibson assembly reaction (Gibson Assembly^®^ Cloning Kit, New England BioLabs) following the manufacturer’s instructions (Figure S4C) (Gibson et al., 2009). The complete ampicillin resistance plasmid was amplified in Electrocomp™ E. coli using a standard protocol (One Shot™ TOP10 Electrocomp™ E.coli, ThermoFisher scientific). The successful assembly of the insertion, REST fragments and the backbone vector, was verified by sequencing, and by PCR where the custom construct was digested with AfIII and AgeI to release REST insert of ~3300bp. The digested products were size-separated by electrophoresis on a 1% agarose gel to check for successful cloning (Figure S5D).

### RNA extraction and quantitative (q)RT-PCR

Cells were lysed with QIAzol Lysis Reagent (Qiagen) and stored at −20°C. Total RNA was extracted from the hiPSCs and hiPSCs-derived neurons lysates using the miRNeasy mini kit (reference 217004, Qiagen, Germany). For each sample, 1μg of total RNA was reverse transcribed using the miScript II RT Kit (Qiagen). For qRT-PCR analysis of miRNAs, the miScript SYBR Green PCR kit (218073; Qiagen, Germany) was employed. While qRT-PCR analysis for other genes was performed using QuantiTect SYBR Green PCR kit (Qiagen, Germany) on a StepOnePlus™ Real-Time PCR System (Applied Biosystems) following the manufacturer’s instructions. All reactions were performed in triplicate for each sample. The relative expression levels of the miRNAs and other genes was calculated using the 2^-ΔΔCT^ method (Livak and Schmittgen, 2001) and the data were normalized to GAPDH and Colrf43. The primer sequences for mature miRNAs as well as all genes examined in the current study are listed in Table S3.

### Western blotting

Culture cells were washed with ice-cold PBS, cells were scraped using cold plastic cell scraper. Cells were treated with ice-cold RIPA buffer (Sigma) and protease inhibitor cocktail (Sigma) and agitated for 30min at 4°C. Cells were centrifuged at 21000 rcf at 4°C and the supernatant was collected. The supernatant was treated with appropriate amounts of LDS sample buffer (NuPAGE) and sample reducing agent (NuPAGE), then samples were heated at 95°C for 5 min. Electrophoresis was carried out using Bolt 4%–12% Bis-Tris Plus Gels (Life Technologies) with 15 μg of protein loaded per sample. Samples were transferred to nitrocellulose membranes and these blots were incubated in blocking solution 5% (w/v) powdered milk in Tris-buffered saline containing 0.1% (v/v) Tween 20 (TBST) for 60 min, at RT. Blots were then incubated overnight at 4°C with primary antibody against REST (1:500) (ab75785, ab21635, Abcam), H3k9me2 (1:500) (ab1220, Abcam), MAP2 (1:750) (MAB8304, R&D Systems, Minneapolis, MN) or Caspease-3 (1:500) (9662, Cell Signaling Technology, USA) diluted in the blocking solution. After washing in TBST, blots were incubated with an appropriate IRDye^®^-conjugated secondary antibody (LI-COR). Proteins were visualised using the Licor/Odyssey infrared imaging system (Biosciences, Biotechnology). Bands were analysed by densitometry using the Odyssey application software and expressed as density of the protein of interest normalised to that of GAPDH.

### Immunocytochemistry

Cultured cells were washed with PBS and fixed in 3.7% PFA for 20 minutes at 4°C. Cells were blocked for 1h in PBS with 0.3% Triton-X-100 (PBS-T) and 5% donkey serum. Cells were incubated with appropriate primary antibodies in PBS-T with 5% donkey serum overnight at 4°C. Secondary antibodies were prepared in PBS-T and left for 1.5h at RT. DAPI (Molecular Probes) was used to counterstain cell nuclei. Staining was preserved using DAKO fluorescent mounting medium (Life Technologies). Samples were imaged on a Leica DMI6000b fluorescent microscope, while cell counts and intensity were determined using CX7 High-Content Screening (HCS) Platform (Thermo Fisher Scientific). Cell counts and intensity from at least 3 replicates were used for statistical analysis. Primary antibodies and their dilutions were used as follows: Nanog (1:200, 4903, Cell Signaling Technology, USA), Oct-4 (1:200, 2750, Cell Signaling Technology, USA), Sox2 (1:200, 3579, Cell Signaling Technology, USA), NeuN (1:250, MAB377, Sigma). Secondary antibodies used were: Alexa 594-conjugated donkey anti-rabbit (1:1000, Invitrogen, A21207), Alexa 488-conjugated donkey anti-rabbit (1:1000, Invitrogen, A21206) and Alexa 488-conjugated donkey anti-mouse (1:1000, Invitrogen, A21202).

### miRNA-seq

The method used to amplify RNA was adapted from Abruzzi et al (Abruzzi et al., 2015) and was performed using TruSeq^®^ Small RNA Library Prep kit (Illumina, USA) and following the manufacturer’s instructions. Briefly, total RNA was extracted from a control hiPSC line and control cells treated with various concentrations of UNC0638 i.e. 50, 100, 200 or 250nM, using the miRNeasy mini kit (Qiagen, Germany). Polyadenylated adaptors were ligated to the 3-end, 5-adaptors were then ligated, and the resulting RNAs were reverse transcribed to generate cDNA that can be amplified by PCR. The amplified product was run on 3% certified™ low range ultra agarose (Bio-Rad Laboratories Ltd) in TBE buffer and a size-selection was performed to ensure that the cDNA used for sequencing primarily contains miRNAs rather than other RNA contaminants. The gel containing the required cDNAs was purified using QIAquick Gel Extraction Kit (Qiagen, Germany). Quality and quantity of miRNA libraries were assessed by Bioanalyzer (Agilent Technologies) and by Qubit (Thermo Fisher Scientific) (Figure S3C). miRNA libraries were then diluted to 2 nM with Illumina RSB buffer and mixed together in equimolar amounts. Libraries were sequenced on an Illumina HiSeq 4000 using single-end 50 base pair reads (10 samples per lane), obtaining a minimum of 35 million mapped reads per sample (Figure S4).

Reads from Illumina single-end sequencing were trimmed with Trimmomatic (Bolger et al., 2014) and assessed for quality using FastQC (https://www.bioinformatics.babraham.ac.uk/projects/fastqc/), using default command line parameters. Reads were mapped to the Homo sapiens GRCh38 reference genome using STAR (Dobin et al., 2012) and counts were assigned to mirbase miRNAs using featureCounts (Liao et al., 2013) and the GRCm38.84 Ensembl gene build GTF. Both the reference genome and GTF were downloaded from the Ensembl FTP site http://www.ensembl.org/info/data/ftp/index.html/. Differential gene expression analyses used the DESeq2 package (Love et al., 2014) and miRNA expression was FPKM-normalised. miRNAs we discarded from the analysis where differential expression failed to be significant between the iPSCs treated with UNC0638 and control untreated condition (significance: adj.pval < 0.05, Benjamini-Hochberg correction for multiple testing). We have also applied thresholds (cutoff) of ≥2.5 fold change in expression.

### miRNA expression analysis

To predict the association between significantly upregulated miRNAs and neurodevelopmental disorders, a crossover analysis between miRNA gene targets and disease associated genes was performed for Intellectual Disability, Schizophrenia and Autism Spectrum Disorder (ASD). Genes associated with Intellectual Disability and ASD (HP:0001249 and HP:0000717 respectively) were shortlisted from the DECIPHER database (DDG2P - V11.2), whilst Schizophrenia associated genes were shortlisted from the GWAS Catalog - EMBL-EBI (V1.0.2). Predicted miRNA target genes were determined using the miRDB database (V6.0). In order to account for miRNA target score, gene crossover probability was calculated for each miRNA in ‘R’ (V4.0.4), using noncentral hypergeometric distribution, with P□<□.01 considered statistically significant. Crossover probability between disease associated miRNAs and REST targeting miRNAs was assessed for each disorder by calculating hypergeometric probability, with P□<□0.05 considered statistically significant.

### Chromatin immunoprecipitation (ChIP)-qRT-PCR

Chromatin immunoprecipitation was performed as previously described (Koch et al., 2007). Briefly, hiPSCs were cultured in Essential 8™ medium and 10^8^ cells were singularised using accutase and collected by centrifugation before being resuspended in serum free media. Formaldehyde (Sigma) was added at a final concentration of 1% and incubated for 10 min at RT. Glycine (Sigma) was then added at a final concentration of 0.125 M and incubated for 5 min at RT. Cells were then washed in ice-cold PBS and centrifuged for 5min at 4°C and the cell pellet was resuspended in cell lysis buffer (10 mM Tris-HCl pH 8.0, 10 mM NaCl, 0.2% Igepal CA-630, 10 mM sodium butyrate, 50 μg/mL PMSF, 1 μg/mL leupeptin). The lysis was incubated for 10 min on ice and then centrifuged for 5min at 4°C to collect the nuclei, which were resuspended in nuclear lysis buffer (NLB 50 mM Tris-HCl pH 8.1, 10 mM EDTA, 1% SDS, 10 mM sodium butyrate, 50 μg/mL PMSF, 1 μg/mL leupeptin) and then diluted with immunoprecipitation dilution buffer. Chromatin was sheared to a fragment size of ~200-600 bp by sonication cycles of 30 sec ON/30 sec OFF with the Bioruptor^®^ PLUS combined with the Minichiller^®^ Water cooler (Cat. No. B02010003) at HIGH power setting. The resulting sheared product was run on 1% gel and compared to that of non-sonicated DNA (Figure S3B). The chromatin was precleared by adding normal rabbit IgG (Merck Millipore). Subsequently, homogeneous protein G-agarose suspension (Roche) was added to the suspension. The resulting chromatin was used to set up the ChIP assay while 270 μL were used as input control. 10μg of di-methylated histone H3 (ab1220, Abcam) (Fang et al., 2012) or immunoglobulin control were incubated with the chromatin overnight at 4°C. Then homogeneous protein G-agarose suspension was added (Roche) to the suspension. The protein G-agarose was spun down and the pellet washed twice with IP wash buffer 1 (20 mM Tris-HCl pH 8.1, 50 mM NaCl, 2 mM EDTA, 1% Triton X-100, 0.01% SDS), once with 750 μL of IP wash buffer 2 (10 mM Tris-HCl pH 8.1, 250 mM LiCl, 1 mM EDTA, 1% IGEPAL CA630, 1% deoxycholic acid) and twice with 10 mM Tris-HCl 1 mM EDTA pH 8.0. The immune complexes were eluted from the beads using IP elution buffer (100 mM NaHCO3, 0.1% SDS). Eluted and input samples were treated with RNase A (MilliporeSigma) and 5 M NaCl before being incubated at 65°°C for 6h. Then samples were treated with proteinase K (ThermoFisher scientific) and incubated at 45°C overnight. The DNA in both ChIP and input samples was purified using QIAquick PCR Purification Kit (28104, Qiagen, Germany), and qRT-PCR was performed. ChIP enrichment was assessed relative to the input DNA in specific genomic regions. The primers sets are listed in Table S4. Amplified material was detected using QuantiTect SYBR Green PCR kit (Qiagen, Germany) on a StepOnePlus™ Real-Time qPCR System (Applied Biosystems).

### Calcium imaging

Cells prepared for calcium imaging assay were maintained in BrainPhys basal for two weeks prior to performing the experiment. Cells were dissociated into single cells with accutase and plated on coverslips at 50000 cells/cm^2^ and left for a week to settle. Neurons were analysed for spontaneous calcium events using Cal-520™ AM (Abcam) and 20% pluronic acid. On the day of the experiment, cells were exposed to media cocktail containing the dye for 1h, then media were removed and cells washed with PBS. Cells were incubated with fresh BrainPhys media for 30min. Calcium imaging was performed using Zeiss Axio Observer inverted microscope (40x objective) and processed using Zeiss Zen software. After identifying the regions of cells, fluorescent images were taken at a rate of 10Hz using constant streaming mode of the camera at a resolution of 1024×1024 pixels. Regions were imaged for 5 minutes per experiment.

### Calcium imaging analysis

Recorded stacks of images were imported into Fuji (Schindelin et al., 2012). Region of interests (ROIs) were then identified in each sub-region using Matlab based NeuroCa package (Jang and Nam, 2015), which adopts an approach based upon circular Hough transforms (Jiang and Ehlers, 2013). ROIs were limited to cell bodies and were detected using a radius threshold of 2-7 pixels. While ROI masks were created with NeuroCa, the analysis of calcium events was performed with the Matlab based FluroSNAAP (Patel et al., 2015). In order to use the ROI mask created with NeuroCa in FluroSNAAP, mask files were converted accordingly using a custom written Matlab script. Files were then processed in FluroSNAAP using the batch processing function to analyse single ROI events only. Results from the analysis were then imported into Excel for processing and subsequently into Prism for statistical evaluation.

### Statistical analysis

Prism 7.0 (GraphPad Software) was used for the statistical analysis. Data shown are the mean□±□SEM. with P□<□.05 considered statistically significant. Two-tailed unpaired t-tests were used for comparisons between two groups. Group differences were analysed with one-way analysis of variance (ANOVA) followed by Tukey’s multiple comparisons test. Data distribution was assumed to be normal, but this was not formally tested.

## Acknowledgment

We thank Dr Janet Harwood for help with bioinformatics. We thank J. Morgan for technical assistance and the DEFINE field team, particularly Rachael Adams, Alister Baird and Dr Stefanie Linden for patient recruitment and assessment (Division of Psychological Medicine and Clinical Neurosciences, Cardiff School of Medicine). We are grateful to the patients and their families for the support of this research. This work was supported through DEFINE, a Wellcome Trust Strategic Award (100202), a Wellcome Trust PhD programme award to BAD (WT093765MA) and The Waterloo Foundation Changing Minds Programme.

